# MLL1 directs gut-associated antibody responses to helminth and bacterial infections

**DOI:** 10.1101/2024.08.21.609083

**Authors:** Yan Zhang, Clarissa Chakma, Alana Kirn, David Chisanga, Jack Polmear, Aidil Zaini, Rohia Farighi, Liang Xie, Diana López-Ureña, Steven Mileto, Dena Lyras, Colby Zaph, Joanna R Groom, Kim L Good-Jacobson

**Affiliations:** Department of Biochemistry and Molecular Biology, Monash University, Clayton, Victoria, Australia 3800; Immunity Program, Biomedicine Discovery Institute, Monash University, Clayton, Victoria, Australia 3800; Department of Immunology, Alfred Medical Research and Education Precinct, Monash University, Melbourne, VIC, Australia; Department of Microbiology, Monash University, Clayton, Victoria, Australia 3800; Infection Program, Biomedicine Discovery Institute, Monash University, Clayton, Victoria, Australia 3800; Division of Immunology, Walter and Eliza Hall Institute of Medical Research, Parkville, Victoria, Australia; Department of Medical Biology, University of Melbourne, Parkville, Victoria, Australia

**Author notes:** Address correspondence to: Kim Good-Jacobson, Department of Biochemistry and Molecular Biology Monash University, Ground Floor Reception, 23 Innovation Walk (Bldg 77) Clayton, Victoria 3800 Australia. Precision Medicine Translational Research Programme, Department of Obstetrics & Gynaecology, Yong Loo Lin School of Medicine, National University of Singapore, Singapore (Liang Xie). Equal contribution (Yan Zhang, Clarissa Chakma).

**Keywords:** B cells, IgA, helminth, trichuris muris, epigenetics, gut-associated lymphoid tissue, MLL1

## Abstract

Soil-transmitted helminths are one of the most common infections globally, yet how to promote effective gut-associated humoral responses is not well understood. We identify the histone methyltransferase MLL1 as a key target to promote IgA-driven responses. *Mll1* was increased in germinal center B cells in gut-associated lymphoid tissues, and *Mll1*-deficiency led to changes in the histone modification H3K4me3 on key B cell and immune-regulatory genes. Correspondingly, MLL1-deficient B cells had defective germinal centers and IgG1 in response to the helminth *Trichuris muris*. Yet, *Mll1*^f/f^*Cd23*^cre/+^ mice expelled worms more rapidly compared to control mice. Accelerated worm clearance correlated with elevated IgA^+^ plasma cells, as well as both serum and fecal IgA. RNA-sequencing identified CCR9 as a key MLL1-regulated molecule. As such, *Mll1*^f/f^*Cd23*^cre/+^ mice infected with *T. muris* had increased IgA^+^CCR9^+^ PC localized in the large intestine. Regulation of IgA by MLL1 was confirmed beyond *T. muris* infection. *In vitro* cultures confirmed *Mll1*-deficiency increased IgA^+^ plasma cells in a B cell-intrinsic manner, and IgA production was also increased in *Mll1*^f/f^*Cd23*^cre/+^ mice infected with the bacterium *C. rodentium*. This study reveals MLL1 as a key target to promote IgA responses to gut-associated infections.

## Introduction

Soil-transmitted helminth infections infect approximately one-quarter of the world’s population, causing severe morbidity and death in high-incidence countries (1–5). Roundworm (*Ascaris lumbricoides*), whipworm (*Trichuris trichiura*) and hookworms (*Necator americanus and Ancylostoma duodenale*) are the most common soil-transmitted helminths that infect humans (6, 7). Currently, the most effective treatment for soil-transmitted helminth infections is to use mass anti-helminthic drug administrations, such as albendazole (8). However, long-term mass anti-helminthic drug therapy induces the development of drug-resistant parasites, and the treatment of reinfection causes a significant economic burden (9–11). Therefore, successful long-term public health interventions such as effective vaccines are urgently needed (12). B cell-mediated antibody responses are the basis of most successful vaccines, yet there is a lack of understanding of how to promote a potent humoral response to intestinal helminth infections.

Helminth-induced immune responses to whipworms such as *T. trichiura* mainly occur in the gut-associated lymphoid tissues (GALT) and are largely dominated by Th2-biased responses (13). Humoral responses in the GALT are highly specialized to maintain gut homeostasis by regulating commensal microbiota and responses to food antigens, pathogenic bacteria or helminths (14). The isotype of antibody produced during infection is critical for the effectiveness of the response (15). While IgG, IgE and IgA are all produced during mucosal responses (8), IgA is particularly crucial for the protection of the human intestinal mucosa (16–18). Increased IgA has been correlated to reduced helminthiasis, increased microbiota diversity and improved responses to pathogenic bacteria such as *Salmonella* Typhimurium (19–25).

Pinpointing key drivers of effective B cell responses in the GALT may aid development of effective mucosal vaccines and new treatments for helminth infections. The epigenetic regulator mixed-lineage leukemia 1 (MLL1; also known as KMT2A) is a histone methyltransferase which promotes B cell development (26). It can also induce *Gata3*, the master transcription factor for Th2 differentiation (27), and regulates Muc2 production by intestinal stem cells, the major intestinal mucin that can provide protection against worms in the colonic epithelium (28). Recent work has revealed that during immune responses, histone modifiers have unique roles in regulating B cell subset formation, B cell migration, the quality and quantity of antibody, and immunopathology (29–35). Given the role of MLL1 in promoting Th2 cells through cooperation with c-Myb (36), a transcription factor we have previously found to be important in tailoring effective antibodies in response to a Th2-induced microenvironment (15), we hypothesized that MLL1 could be a key regulator of B cell responses in the GALT. Here, we find that conditional deletion of MLL1 in mature B cells promotes IgA responses to the helminth infection *Trichuris muris*. These results identify MLL1 as a therapeutic target to promote the production of IgA^+^ PC.

## Materials and Methods

### Mice

*Cd23*^cre/+^ mice (37) and *Mll1*^f/f^ mice (38) were generously provided by Meinrad Busslinger (Institute of Molecular Pathology, Austria) and Patricia Ernst (University of Colorado, USA), respectively. *Mll1*^f/f^*Cd23*^cre/+^ on C57Bl/6 background were generated by crossing *Cd23*^cre/+^ with *Mll1*^f/f^ mice. Mice were bred at the Monash Animal Research Platform under specific-pathogen-free conditions. All animal experimentation was performed following the Australian National Health and Medical Research Council (NHMRC) Code for the Care and Use of Animals for Scientific Purposes. Procedures were approved by Monash and WEHI Animal Ethics Committees. Both males and females, over 6 weeks old, were used in this study.

*Immunization:* Mice were intraperitoneally immunized with 100μg NP13-KLH precipitated in the adjuvant alum. *T. muris infection:* mice were infected with 200 *T. muris* eggs (39) by oral gavage.

*C. rodentium infection:* mice infected with 2×10^9^ colony-forming units of *C. rodentium* by oral gavage.

### Flow cytometry, cell sorting and antibodies

Single-cell suspensions were stained with fluorochrome-labelled antibodies in PBS 2%FCS for flow cytometric analysis or cell sorting as previously described (35). Rat IgG2b kappa (2.4G2) was used to block the binding of antibodies by cell-surface Fc-receptors and normal rat serum was used to block non-specific binding. For IgA staining, a transcription factor buffer set (BD Pharmingen) was used following the manufacturer’s recommendations. Cells were analyzed using a BD-LSRFortessa X-20 flow cytometer and FACSDiva software (BD Biosciences). Data was analyzed using FlowJo^TM^ software (BD Life Sciences). For cell sorting, cells were sort-purified using a BD Influx cell sorter (BD Biosciences). Antibodies used were as follows. Purchased from Biolegend: CD19 APC-Cy7, CD38 Pacific Blue, IgD AF488, B220 Pacific Blue, IgM PE, IgD PerCP-Cy5.5, CD23 AF488, CD21/CD35 APC, CD19 PE, CD138 PE. Purchased from BD Biosciences: CD95 PE-Cy7, IgG1 APC, CD5-biotin, CD3e APC-Cy7.

### RT-qPCR

Either RNeasy Plus Micro Kit (QIAGEN) or Direct-zol RNA Microprep kit (Zymo) were used to extract RNA from sort-purified cells following the manufacturer’s recommendations, and TRIzol was used to extract RNA from homogenized proximal colon tissue. Reverse Transcription was performed using High-Capacity cDNA Reverse Transcription kit (Thermo Fisher Scientific). SYBR Green PCR kit (QIAGEN) was used to perform qPCR. *Rpl32* was used as an internal control for normalization and calculating the ΔΔCt value (40). Primers used were: *Mll1* 5ʹ-GCAGATTGTAAGACGGCGAG-3ʹ, 5ʹ-GCAGATTGTAAGACGGCGAG-3ʹ; *Rpl32* 5ʹ-ATCAGGCACCAGTCAGACCGAT-3ʹ, 5ʹ-GTTGCTCCCATAACCGATGTTGG-3ʹ.

### Worm counting

To determine worm clearance, ceca were collected from *T. muris*-infected mice, worms extracted and counted.

### T. muris-specific ELISA

High-binding 96-well ELISA-plates (Sarstedt) were coated with *T. muris* antigen (39) diluted in carbonate buffer, and incubated overnight at 4℃. Following wash and blocking steps, samples that were serially diluted and incubated for 1 hour at room temperature. After washing, anti-IgG1-HRP (Southern Biotech) was added and incubated for 45 minutes at room temperature. The wells were washed and TMB substrate (eBioscience) added, and after 1 minute, stopped with 1N HCl. Plates were read at 450nm using a BMG OPTIMA plate reader.

### Serum IgA ELISA and fecal IgA ELISA

*Sera:* blood was collected by cardiac puncture immediately post-mortem and sera collected post-centrifugation. *Fecal extraction:* performed as previously described (41, 42) by adding 0.1g of fecal pellets to 1ml of PBS containing complete mini protease inhibitor cocktail (Roche). Solids were separated by centrifugation and supernatants (fecal extract) stored at -20℃ until use. *IgA ELISAs*: high-binding 96-well ELISA plates were coated using unlabelled IgA coating antibodies (Southern Biotech). Following wash and blocking steps, samples that were serially diluted and incubated for 37℃ for 3 hours. Following washing, anti-IgA-HRP (Southern Biotech) was added and incubated for 1 hour at 37℃. Plates were washed and developed with OPD-substrate solution (Sigma-Aldrich). The plates were read at 450nm using a BMG OPTIMA plate reader.

### Immunohistochemistry

mLN from *T. muris*-infected mice were first fixed in periodate-lysine-paraformaldehyde fixative, immersed in 30% sucrose and then subsequently embedded and frozen in Optimal Cutting Temperature compound (Tissue-Tek). 7μm tissue sections were cut using a microtome (Leica) and mounted on Superfrost Plus slides (Thermo Fisher). Sections were fixed with cold acetone (Sigma) at -20°C for 10 minutes. Sections were stained with PNA (Vector Labs) and B220 (BD Bioscience). Slides were observed and images were captured under 4X and 10X magnification using a Nikon microscope.

### In vitro B cell culture

B cells from the spleen were isolated using the EasySep™ Mouse B Cell Isolation Kit (STEMCELL Technologies) and labeled with Cell Trace Violet (ThermoFisher). 5×10^4^ labeled cells were resuspended in B cell media (RPMI, 5%FCS, 50μM 2-mercaptoethanol and 2mM glutamine) and stimulated with 50ng/ml CD40L (R&D Systems), 50ng/ml IL-4 (STEMCELL Technologies), 5ng/ml IL-5 (R&D Systems), 10ng/ml TGFβ (R&D Systems) and 10nM retinoic acid.

### RNA-sequencing

Germinal center (GC) B cells (B220^+^IgD^lo^CD95^hi^CD38^lo^) were sort-purified from mLN of *T.muris*-infected *Cd23*^cre/+^ and *Mll1*^f/f^*Cd23*^cre/+^ mice. RNA was extracted using the RNeasy Plus Micro Kit according to the manufacturer’s instructions. Total mRNA was quantified with the Qubit Fluorometer (Invitrogen) and RNA integrity number values were assessed using the Agilent Bioanalyzer. mRNA libraries were sequenced on an Illumina Novaseq using a 2x150 paired-end (PE) configuration according to the manufacturer’s instructions. Data was processed and analysed as previously described (33, 35). The following values were used to detect differentially expressed genes: fold change >2; q-value (FDR-adjusted p-value) <0.05. KEGG (Kyoto Encyclopedia of Genes and Genomes) enrichment analysis was performed using KEGG databases. Data has been deposited at GEO and publicly available. Accession number is GSE237391.

### CUT&Tag sequencing

Cleavage Under Targets and Tagmentation (CUT&Tag) was performed for H3K4me3 on sort-purified mesenteric lymph node GC B cells (B220^+^IgD^lo^CD95^hi^ CD38^lo^CD138^-^) from *T. muris*-infected *Mll*^f/f^*Cd23*^Cre/+^ and *Mll*^f/f^*Cd23*^+/+^ mice at day 14 post-infection based on the original CUT&Tag protocol described previously (43), with some variations. Nuclei were extracted by resuspending cells in cold nuclear extraction buffer (20mM HEPES [pH 7.9], 10mM KCl, 0.1% Triton X, 10% glycerol, 0.5mM spermidine) with cOmplete™, EDTA-free Protease Inhibitor (Roche). Nuclei were spun down and resuspended in 100μl of nuclear extraction buffer, before incubated with 11μl of activated concanavalin A-coated magnetic beads (Bangs Laboratories) for 10 minutes at room temperature. Bead-bound nuclei were resuspended in 50μl of Dig150 buffer (20mM HEPES [pH 7.9], 150 mM NaCl, 0.5mM spermidine, 0.01% digitonin) containing 0.5μg of H3K4me3 antibody (Cell Signalling Technology) and incubated overnight at 4°C.

Primary antibody solution was discarded and bead-bound nuclei were resuspended in 50μl of Dig150 buffer containing 0.5μg of secondary antibody (Antibodies Online) and incubated for 30 minutes at room temperature. Following incubation, bead-bound nuclei were washed gently with Dig150 buffer. Bead-bound nuclei were resuspended with 50μl of Dig300 buffer (20mM HEPES [pH 7.9], 300 mM NaCl, 0.5mM spermidine, 0.01% digitonin) containing pAG-Tn5 (EpiCypher) for 1 hour at room temperature, with gentle agitation/shaking. Following incubation, bead-bound nuclei were washed gently with Dig300 buffer and resuspended in 50μl of tagmentation buffer (Dig300 buffer with 10mM MgCl2) and incubated for 1 hour at 37°C. To each reaction, 50μl of TAPS buffer (10mM TAPS [pH 8.5], 0.2mM EDTA) and 25μl of AMPure beads (Beckman Coulter) was added and mixed thoroughly, before adding 7μl of SDS Release Buffer (TAPS buffer with 0.1% SDS) to release Tn5. Samples were incubated for 1 hour at 58°C in a thermocycler (heated lid at 95°C). To neutralise the SDS, 10μl of 1% Triton-X 100 was added to each sample. Beads were washed with 80% ethanol twice using a magnetic stand and DNA was eluted in nuclease free water.

PCR amplification of CUT&Tag libraries was performed by mixing 25μl of NEBNext® High-Fidelity 2X PCR Master Mix (New England Biolabs), 22μl of tagmented DNA, 2μl of uniquely barcoded i5 (at 10μM) and 2 μl of uniquely barcoded i7 primer (at 10μM). PCR was performed as follows: 58°C for 5 min; 72°C for 5 min; 98°C for 45 sec; 16 cycles of 98°C for 15 sec and 63°C for 10 sec; a final extension at 72°C for 1 min and hold at 4°C. Libraries were purified post-PCR by mixing 65μl of AMPure beads and incubating for 10 minutes at room temperature, followed by two washes with 80% ethanol using a magnetic stand. Sequencing libraries were eluted in 10mM Tris-HCl (pH 8). Library concentrations and size distributions were assessed by Qubit and Bioanalyzer. Libraries were pooled at an equimolar ratio and sequenced on the Nextseq2000 using 100 cycle chemistry.

### CUT&Tag sequencing analysis

Processing of the raw data, including quality control, adapter trimming, alignment to the mouse reference genome (mm10), and generation of intermediate bigwig files, was conducted using nf-core/cutandrun (V3.2.2) (44), a Nextflow-based pipeline. Enriched H3K4me3 associated histone peaks were identified using MACS3 (45) with an FDR cut-off of 5%. A consensus set of peaks was determined and annotated to various genome regions using Homer’s annotatePeaks function (46). The summarisation of read counts for the consensus peaks was achieved using the featureCounts (47) function within Rsubread (48). The limma-voom pipeline (49) was used to identify differentially accessible regions. Briefly, peaks with a count per million (cpm) < 5 in each of the libraries were excluded from subsequent analysis. Read counts were converted to log2(cpm), quantile normalised, and precision weighted with the voom function of the limma package (50). A linear model was fitted to each peak, and empirical Bayes’ moderated t-statistics utilised to assess differences in peak occupancy. Peaks were called as differentially accessible if they achieved a false discovery rate (FDR) <0.1. The enrichment of Gene Ontology terms and KEGG pathways was performed using the goanna and kegga functions within limma.

### Statistical analysis

Experimental data are presented as mean ± standard error of the mean (SEM). Statistical analyses performed with Mann-Whitney non-parametric test using GraphPad Prism 8 (GraphPad Software).

\**p* < 0.05, \*\**p* < 0.01 and \*\*\**p* < 0.001. Mice assessed to be non-responders were excluded from all analyses.

## Results

### MLL1 is required for optimal germinal center B cell responses

To investigate whether MLL1 may tailor mucosal B cell responses, we first examined the expression of *Mll1* in mature B cell subsets in GALT. B cell subsets were sort-purified from naïve wild-type mice and assessed for *Mll1* expression by RT-qPCR in isolated mesenteric lymph nodes (mLN) and Peyer’s patches. At steady-state, there is an active immune response to microbiota in the gut-draining secondary lymphoid organs, in which activated B cells can form extrafollicular antibody-secreting PC, or GC (2, 51). GCs provide a site for affinity maturation of B cells, enabling the production of long-lived memory B cells and PC, two populations that can persist for long periods of time to provide immunity (52). In comparison to naïve B cells and PC, GC B cells had significantly increased expression of *Mll1* (Fig. 1A-B). To determine whether this increase in *Mll1* in GC B cells was observed systemically, we immunized wild-type mice with (4-hydroxy-3-nitrophenyl)acetyl conjugated to keyhole limpet hemocyanin (NP-KLH), a haptenated protein precipitated in the adjuvant alum, and assessed *Mll1* in sort-purified subsets post-immunization. Commensurate with GALT GCs, *Mll1* was significantly upregulated in splenic GC B cells compared to follicular B cells (Fig. 1C). A small but non-significant increase in *Mll1* expression was also detected in memory B cells, compared to follicular B cells (Fig. 1C).

**Figure 1.**
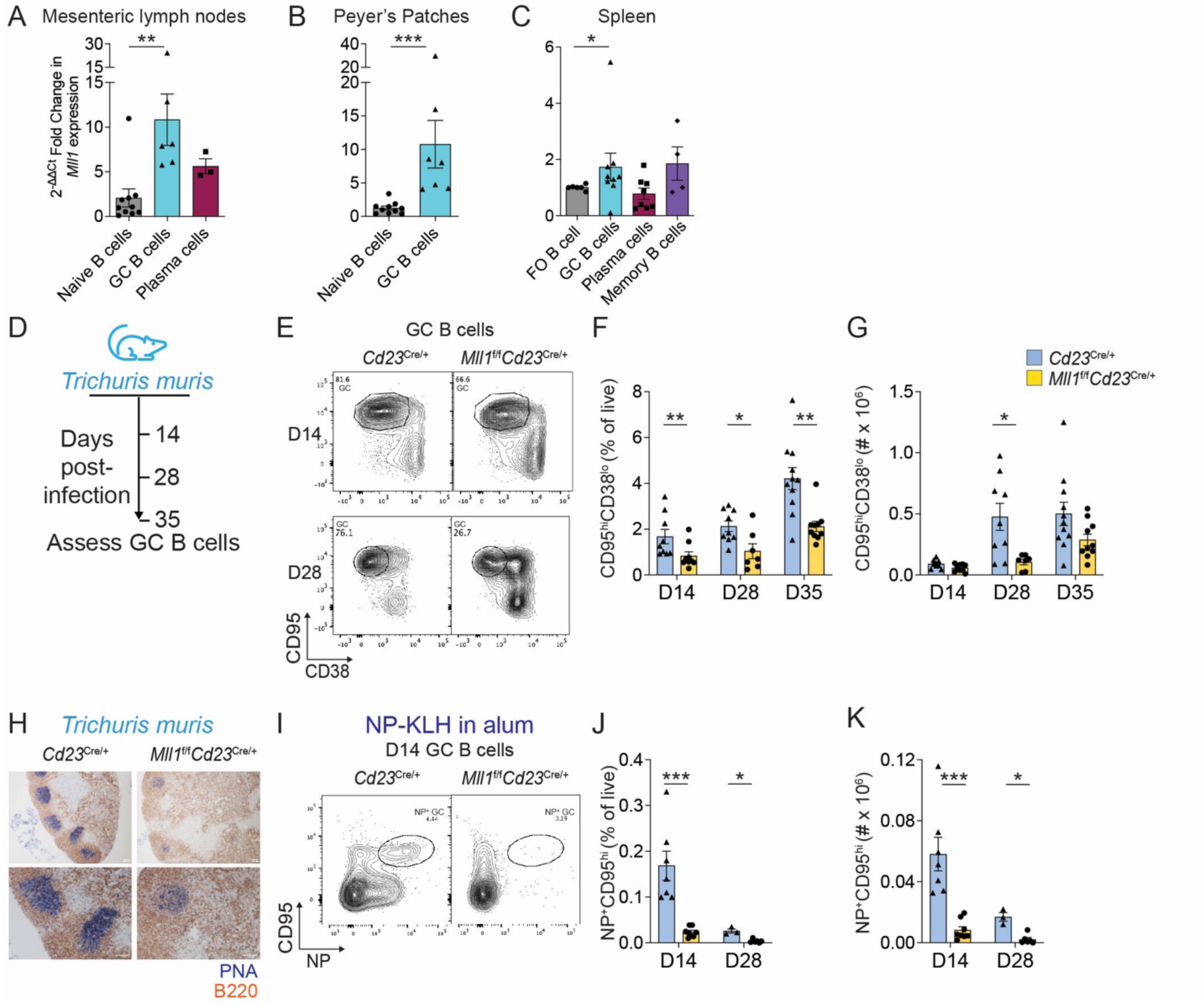
MLL1 is required for GC B cell differentiation in Th2 cell-biased responses. (A and B) The expression of *Mll1* was assessed by RT-qPCR in B cell subsets sort-purified from (A) the mesenteric lymph node (mLN) and (B) Peyer’s patches of C57Bl/6 mice. (C) Follicular (FO) B cells, GC B cells, PC and memory B cells were sort-purified from spleen and *Mll1* expression assessed. FO B cells were sort-purified from C57Bl/6 mice, GC B cells were sort-purified from C57Bl/6 mice immunized with NP-KLH in alum at d7 post-immunization, and memory B cells and PC were sort-purified from C57Bl/6 mice immunized with NP-KLH in alum at d28 post-immunization. Expression of *Mll1* was normalized to *Rpl32* and then normalized to naïve (A, B) or follicular (C) B cells. (D) Schematic depiction of *T. muris* experimental setup. (E) Flow cytometric analyses of B220^+^IgD^lo^CD95^hi^CD38^lo^ (GC B cells) in mLN *T. muris*-infected *Cd23*^cre/+^ and *Mll1*^f/f^*Cd23*^cre/+^ mice at d14 and d28 post-infection. The number in plots shows the frequency of GC B cells within the B220^+^IgD^lo^ population. (F and G) Frequency (F) and number (G) of GC B cells post-infection. Data are combined from three independent experiments per time point; n=9 (d14), n=7-9 (d28) and n=10-11 (d35) per genotype. (H) Representative histological images of mLN from d14 *T. muris*-infected *Cd23*^cre/+^ and *Mll1*^f/f^*Cd23*^cre/+^ mice: B220 (red) and PNA (blue). Scale bar = 100μm (I) Flow cytometric analysis of NP^+^ GC B cells (B220^+^IgD^lo^CD95^hi^NP^+^) from d14 post-immunized *Cd23*^cre/+^ and *Mll1*^f/f^*Cd23*^cre/+^ mice. (J and K) Frequency (J) and number (K) of GC B cells post-immunization at d14 and d28. Data are combined from two independent experiments per time point; n=7 (d14) and n=3-6 (d28) per genotype. Error bars indicate mean ± SEM, *p < 0.05, **p < 0.01, ***p < 0.001, Mann-Whitney test.

Given the increased expression of *Mll1* in GC B cells, we assessed whether MLL1 was required for GC formation. To conditionally delete *Mll1* in B cells, we crossed *Mll1* floxed mice to mice in which *Cre* is under the control of the B cell-specific *Cd23* promoter (37). Thus, upon Cre expression, *Mll1* would be expressed normally during B cell development, but conditionally deleted within mature B cells. To assess Th2 responses, we used the *T. muris* model of human *T. trichiura* whipworm infection. *T. muris* is antigenically similar to *T. trichiura*, and both elicit a similar type of immune response, making *T. muris* an effective model to study the B cell response to intestinal helminth infection (13, 53, 54). *Mll1*^f/f^*Cd23*^cre/+^ and *Cd23*^cre/+^ controls were infected with a high dose of *T. muris* to induce a Th2 cell-biased response and B cell differentiation assessed over time (Fig. 1D). The frequency of GC B cells was significantly decreased in *Mll1*^f/f^*Cd23*^cre/+^ mice compared to control mice at each time point post-infection. A 1.5-fold, 4.5-fold and 1.7-fold reduction in the number of GC B cells was observed in B cell-specific MLL1-deleted mice compared to control mice on days (d)14, d28 and d35, respectively (Fig. 1E-G). Histological analyses confirmed that GC structures were less frequent in mLN in the absence of *Mll1* (Fig. 1H).

To understand whether the role for MLL1 in GC biology was limited to GALT, or whether it was generalizable to other types of B cell responses, *Mll1*^f/f^*Cd23*^cre/+^ and *Cd23*^cre/+^ mice were immunized with alum-precipitated NP-KLH and GC B cell formation assessed in the spleen (Fig. I-K). Use of this model antigen allows the assessment of T-dependent B cell differentiation in the absence of the pathogen-influenced changes to the tissue microenvironment that occurs upon infection. In the absence of *Mll1*, GC B cells were virtually absent on d14 and d28 (Fig. 1I-K). Therefore, MLL1 was required for effective GC B cell responses in response to infection or immunization.

### Mll1 deletion induces preferential production of IgA over IgG1

B cells and helminth-specific antibodies can contribute to relieving worm burden (51, 54). Thus, given the reduction of GC B cells, we hypothesized that there would be a reduced ability of MLL1-deficient mice to expel worms. However, worm clearance occurred more rapidly in *Mll1*^f/f^*Cd23*^cre/+^ mice, compared to *Cd23*^cre/+^ controls, with a >3-fold reduction on d14 post-*T. muris* infection (Fig. 2A). Thus, conditional deletion of *Mll1* within mature B cells led to a more efficient and accelerated worm expulsion during the early stages of *T. muris* infection. IgG1 production was next assessed on 14 days post-infection to investigate whether accelerated worm expulsion in *Mll1*^f/f^*Cd23*^Cre/+^ mice correlated to a change in the Ig subclass dominating the response. *T. muris*-specific IgG1 was reduced significantly in the absence of functional MLL1 (Fig. 2B), as was the frequency of IgG1^+^ GC B cells and IgG1^+^ PC (Fig. 2C-E). Thus, these results implied that IgG1 was not responsible for the accelerated worm clearance observed in *Mll1*-deficient mice. Apart from antigen-specific antibodies, Th2 cytokines such as IL-13 and IL-4 are known to limit worm burden in human or murine intestinal helminth infection (55, 56). To examine the possibility that accelerated worm clearance may be downstream of a functional change in cytokine production, cytokines were assessed in homogenized proximal colon tissue of d14 *T. muris*-infected mice by qPCR. However, no significant change was detectable (data not shown). We also assessed the frequencies of Th2 and T regulatory cell subsets, which were found to be unchanged between *Mll1*^f/f^*Cd23*^Cre/+^ and *Cd23*^Cre/+^ mice (data not shown).

**Figure 2.**
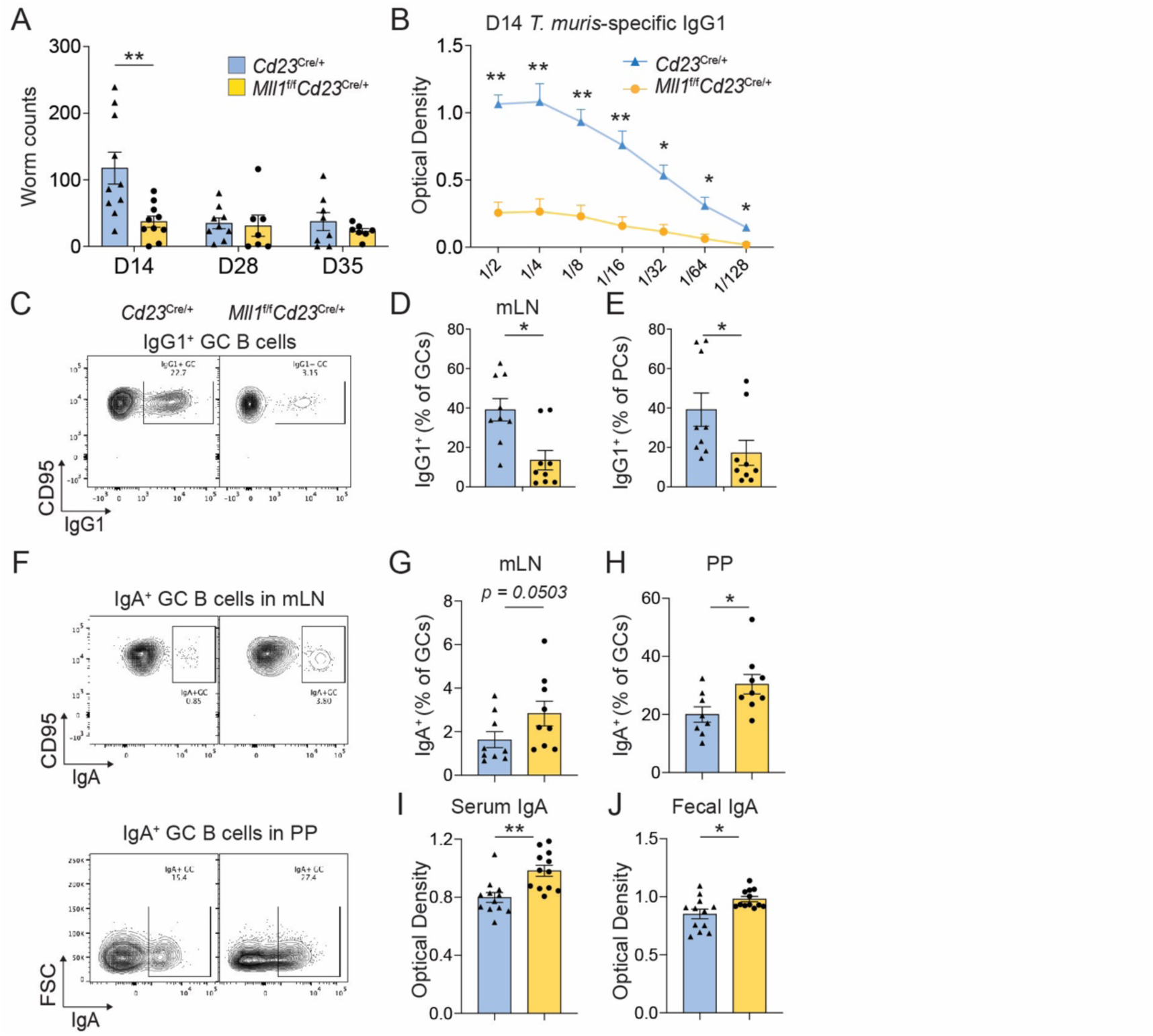
B cell-intrinsic MLL1 deletion leads to rapid worm clearance and increased IgA. (A) Number of worms in the caecum from *T. muris*-infected *Cd23*^cre/+^ and *Mll1*^f/f^*Cd23*^cre/+^ mice at d14, d28 and d35 post-infection. Data are combined from 2-3 experiments per time point; n=10 (d14); n=7-9 (d28); n=7-8 (d35) per genotype. (B) *T. muris*-specific serum IgG1 at d14 post-infection. (C) Representative flow cytometric plots of IgG1^+^GC B cells (IgG1^+^CD95^hi^CD38^lo^) in mLN at d14. The number in plots shows the frequency of IgG1^+^ within the GC B cell population. (D-E) Frequency of (D) IgG1^+^ GC B cells and (E) IgG1^+^ PC (IgG1^+^B220^lo^CD138^hi^). Data are combined from three independent experiments, n=9 per genotype. (F) Representative plots shown of IgA^+^ GC B cells (IgA^+^CD95^hi^CD38^lo^) in mLN or Peyer’s patches at d14. The number in plots shows the frequency of IgA^+^ within the GC B cell population. (G and H) Frequency of IgA^+^ GC B cells in the (G) mLN and (H) Peyer’s patches. Data are combined from two independent experiments. n=8 or 9 per genotype. (I) Serum IgA and (J) fecal IgA from d14 *T. muris*-infected *Cd23*^cre/+^ and *Mll1*^f/f^*Cd23*^cre/+^ mice. Data are combined from 3-4 independent experiments, n=12 per genotype. Error bars indicate mean ± SEM. *p < 0.05, **p < 0.01, Mann-Whitney test.

IgA is also key in the protection of human and murine intestinal mucosa (16, 18). IgA antibodies have been linked to *T. muris* worm expulsion from the intestine (25). Thus, we investigated whether MLL1 regulates the production of IgA. As mLN and Peyer’s patches are both IgA-inductive sites in the intestinal system (57), IgA^+^ cells were studied in both sites on d14 post-infection. Both mLN and Peyer’s patches showed increased IgA^+^ GC after B cell-intrinsic deletion of *Mll1* (Fig. 2F-H). Accordingly, serum total IgA (Fig. 2I) and fecal total IgA (Fig. 2J) were both significantly increased in *T. muris*-infected *Mll1*^f/f^*Cd23*^Cre/+^ mice compared to control mice. It was possible that this increase in IgA may already have been present in *Mll1*^f/f^*Cd23*^Cre/+^ mice prior to infection, and not directly driven by the B cell response to *T. muris* infection. We therefore assessed IgA levels in uninfected mice. Both fecal total IgA and serum total IgA were comparable between uninfected *Cd23*^cre/+^ and *Mll1*^f/f^*Cd23*^Cre/+^ mice (data not shown), indicating that the elevated IgA detected in the absence of *Mll1* was specifically induced by *T. muris* infection. Thus, while IgG1^+^ cells were decreased in the absence of MLL1 *in vivo*, IgA was elevated in response to infection and correlated with increased pathogen clearance.

To examine whether the increased production of IgA observed *in vivo* was due to B cell-intrinsic or B cell-extrinsic mechanisms, we assessed the ability of MLL1-deficient B cells to differentiate into IgA^+^ PC *in vitro*. Naïve B cells from *Mll1*^f/f^*Cd23*^cre/+^ and *Cd23*^cre/+^ mice were stimulated *in vitro* with IgA-promoting stimuli (CD40L, IL-4, IL-5, TGFβ and retinoic acid). After 4 days, plasmablasts were assessed. MLL1-deficient cells had a significantly increased propensity to form IgA^+^ plasmablasts, compared to control B cells (Fig. 3A-C). This was also reflected in the increase in IgA secreted in the cultures (Fig. 3D). Therefore, in the absence of MLL1, B cells were intrinsically prone to produce IgA-producing PC. Taken together, the data thus far demonstrated that MLL1 regulated IgA production in B cells.

**Figure 3:**
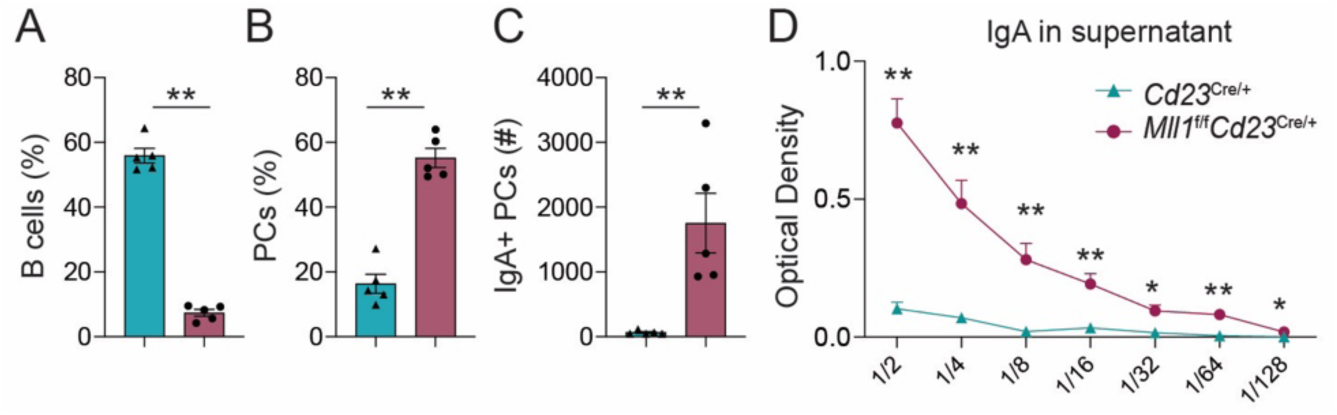
Increase in the production of IgA^+^ PC *in vitro* in the absence of *Mll1*. B cells were stimulated in culture with CD40L, IL-4, IL-5, TGFβ and retinoic acid for 4 days. (A) Frequency of B cells, (B) frequency of PC, (C) total number of IgA^+^ PC, and (D) IgA in supernatant at d4 post-stimulation. Data are combined from two individual experiments, n=5 per genotype. Error bars indicate mean +-SEM, *p < 0.05, **p < 0.01, Mann-Whitney test.

### MLL1 regulates intestinal localization of IgA^+^CCR9^+^ PCs

We next investigated the transcriptional program altered in the absence of MLL1 that may tailor B cell responses in GALT. GC B cells (CD19^+^IgD^lo^CD95^hi^CD38^lo^) were sort-purified from mLN of *T. muris*-infected *Cd23*^cre/+^ and *Mll1*^f/f^*Cd23*^cre/*+*^ mice and RNA-sequencing undertaken. 34 genes with a >2-fold expression change was observed in GC B cells after B cell-intrinsic deletion, including 7 down-regulated genes and 27 up-regulated genes (Fig. 4A). Differential gene KEGG enrichment analysis was performed to get further insight into the gene pathway in GC B cells mediated by conditional deletion of *Mll1* in B cells. Consistent with the increased IgA in MLL1-deficient mice, KEGG analysis identified a gene signature related to ‘an intestinal immune network for IgA production’, along with several disease and stress pathways (Fig. 4B). Specifically, this analysis identified *Ccr9* as significantly upregulated in *Mll1*^f/f^*Cd23*^cre/*+*^ mice than *Cd23*^cre/+^ mice (Fig. 4C).

**Figure 4.**
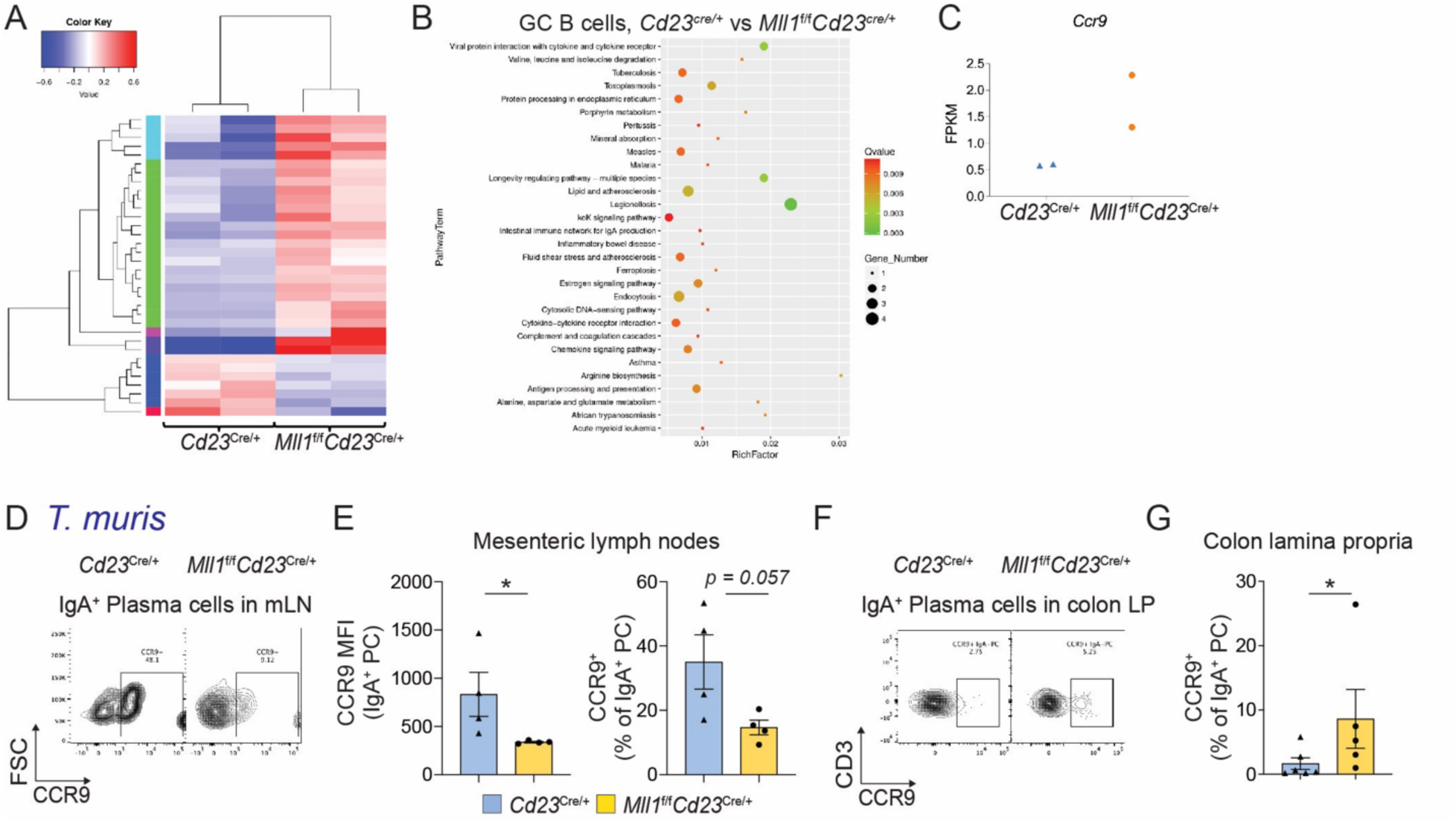
MLL1 regulates the expression of genes related to IgA production. (A) Heatmap of RNA-sequencing data with fold change >2 and q-value (FDR-adjusted p-value) < 0.05; each column is an independent sample obtained from sort-purified GC B cells (B220^+^IgD^lo^CD95^hi^CD38^lo^) from mLN of d13 *T.muris*-infected *Cd23*^cre/+^ and *Mll1*^f/f^*Cd23*^cre/+^ mice, n=2 per genotype. (B) KEGG enrichment pathway analysis of differential gene expression. The size of the dot is positively correlated with the number of differential genes in the pathway. (C) Plots showing FPKM (Fragments per kilo base of transcript per million mapped fragments) for *Ccr9*. (D) Representative flow cytometric plots of CCR9^+^IgA^+^ PC (CCR9^+^IgA^+^B220^lo^CD138^hi^) in mLN from d14 *T. muris*-infected *Cd23*^cre/+^ and *Mll1*^f/f^*Cd23*^cre/+^ mice. (E) CCR9 geometric mean fluorescence intensity (MFI) and CCR9^+^ frequency within IgA^+^ PC in mLN; n=4 per genotype. (F) Representative plots of CCR9^+^IgA^+^ PC (CCR9^+^IgA^+^CD3^-^B220^lo^Blimp1^+^) in colon lamina propria from d14 *T. muris*-infected *Cd23*^cre/+^ and *Mll1*^f/f^*Cd23*^cre/+^ mice. (G) Frequency of CCR9^+^ cells in IgA^+^ PC population in colon lamina propria. Data shown are from 2 experiments, n=6 per genotype. Error bars indicate mean ± SEM. *p < 0.05, Mann-Whitney test.

Upregulation of CCR9 induces the migration of IgA^+^ PCs from mLN to the intestinal lamina propria in response to CCL25 (58). Given that *Ccr9* was identified as differentially expressed in the absence of *Mll1*, we examined whether MLL1 regulated CCR9 expression on IgA^+^ PCs in GALT. In contrast to GC B cells (Fig. 4A-C), the frequency of CCR9^+^ IgA^+^ PCs and CCR9 expression on IgA^+^ PCs in mLN were decreased in MLL1-deficient mice (Fig. 4D-E), whereas there was no change detected in the small intestine (data not shown). As *T. muris* infection mainly occurs in the large intestine, including colon (59), we hypothesized that CCR9 may be regulating IgA^+^ PC migration to the colon lamina propria (Fig. 4F). Indeed, the frequency of CCR9^+^ IgA^+^ PCs was significantly increased in colon lamina propria of *Mll1*^f/f^*Cd23*^cre/*+*^ mice compared to *Cd23*^cre/+^ mice, respectively (Fig. 4G). Thus, these results indicated that B cell-specific *Mll1* deletion increased CCR9^+^IgA^+^ PCs to the colon.

To confirm that the observed regulation of IgA PC by MLL1 was not unique to the *T. muris*-induced GALT immune response, we used the bacteria *Citrobacter rodentium*, which also induces IgA production (60). *Mll1*^f/f^*Cd23*^cre/+^ and *Cd23*^cre/+^ mice were infected with *C. rodentium* and responses assessed at d14 post-infection (Fig. 5A). In accordance with the humoral response to *T. muris*, *Mll1-*deficiency resulted in an increase in serum IgA (Fig. 5B). Taken together, the absence of *Mll1* in B cells leads to increased production of IgA^+^ PC *in vitro*, localization of CCR9^+^ IgA^+^ PCs in the colon lamina propria in response to *T. muris,* and increased IgA production post-infection with either *T. muris* or *C. rodentium in vivo*.

**Figure 5:**
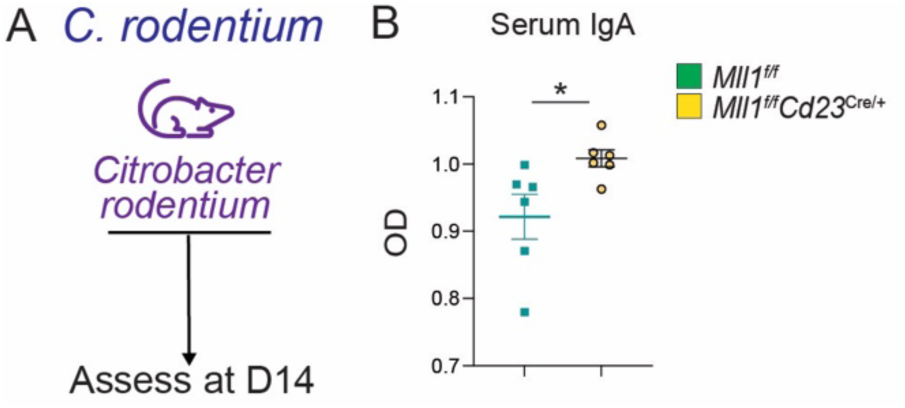
MLL1 regulates B cell responses to *C. rodentium*. (A) *Mll1*^f/f^ and *Mll1*^f/f^*Cd23*^cre/+^ mice were infected with *C. rodentium* and assessed at d14 post-infection. (B) Serum IgA, combined from two independent experiments. *p < 0.05, Mann-Whitney test.

### MLL1 modulates H3K4me3 peaks at immune regulatory gene loci

Lastly, we assessed the role of MLL1 in regulating H3K4me3 within mucosal B cells. Technical limitations precluded deep-sequencing of the PC population, therefore we proceeded to investigate histone modifications within GC B cells in the GALT. Flow cytometric and histological analyses (Fig. 1) demonstrated an abrogation of GC B formation in *Mll1*^f/f^*Cd23*^cre/+^ mice. This was opposite to that of KMT2D (MLL2), in which deletion of *Kmt2d* during B cell development promoted proliferation and GC B cell formation (31). Both histone modifiers have been shown to regulate H3K4me3, a histone modification that is associated with euchromatin and thus promotion of gene expression (61). We therefore tested whether MLL1 regulated H3K4me3 in mLN GC B cells post-infection with *T. muris*. To do this, we undertook CUT&Tag sequencing, which enables assessment of histone modifications at site-specific loci (43). The majority of H3K4me3 peaks mapped to either the promoter-TSS sites (30.5%) or intronic regions (37.6%; Fig. 6A). In contrast to *Kmt2d* deletion, in which H3K4 methylation was globally reduced (31), alterations in H3K4me3 in *Mll1*-deficient GC B cells appeared to be more selective and were either increased or decreased, depending on the gene loci (Fig. 6B-E). Differential peak analyses revealed that 296 regions were altered in the absence of MLL1, with 112 peaks upregulated and 184 peaks downregulated in *Mll1*^f/f^*Cd23*^cre/+^ mice, compared to *Cd23*^cre/+^ mice (Fig. 6B). Concomitant with known MLL1 function, Gene Ontology and KEGG pathway analysis demonstrated that the top 20 molecular functions of genes associated with differentially detected peaks were mainly DNA-binding and nucleic acid-binding activity, along with growth factor, coreceptor and receptor ligand activity (Fig. 6C). We then interrogated the differential peak analyses further to determine whether any genes associated with B cell biology, or genes that were identified as differentially expressed by the RNA-sequencing (Fig. 4). We identified several loci at which H3K4me3 was modulated by the absence of MLL1 (Fig. 6D-E), including migratory and/or B cell factors *Cxcl12, Cd72, Cd74, Klf9, Tbx21 and Fcer2a* (Fig. 6D). Several genes that were differentially expressed (Fig. 4) were confirmed to be direct targets of MLL1-mediated regulation of H3K4me3, such as *Dkk3, Gng3, and Cfp* (Fig. 6E). Therefore, while MLL1 deficiency did not lead to a global absence of H3K4me3, it was involved in fine-tuning H3K4me3 at immune-regulatory gene loci during mucosal immune responses.

**Figure 6:**
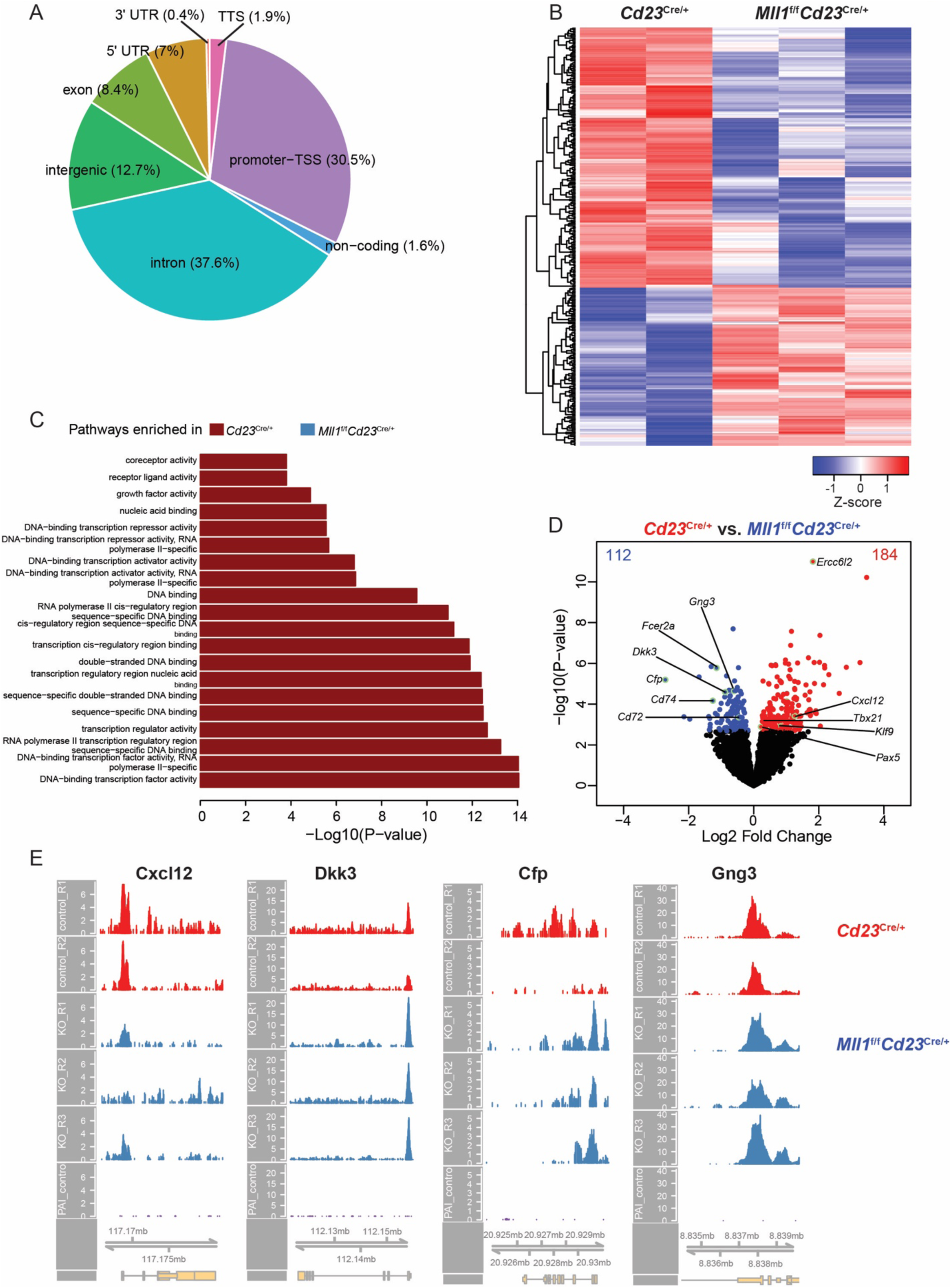
*Mll1*-deficiency induces H3K4me3 alterations at key gene loci. (A) Pie chart illustrating the proportion of peaks annotated to various regions in the genome. (B) Heatmap displaying all differentially accessible regions between *Cd23*^cre/+^ and *Mll1*^f/f^*Cd23*^cre/+^ mice. (C) Bar plot of enriched molecular function Gene Ontology terms for DARs between *Cd23*^cre/+^ and *Mll1*^f/f^*Cd23*^cre/+^ mice. (D) Volcano plot depicting DARs between *Cd23*^cre/+^ and *Mll1*^f/f^*Cd23*^cre/+^ mice. (E) Coverage plots illustrating chromatin accessibility in the *Cxcl12*, *Dkk3*, *Cfp*, and *Gng3* gene regions, extending 3000bp before the start and 1000bp at the end of the gene body.

## Discussion

Uncovering key regulators of mucosal immune responses is important for designing vaccines that can induce effective protective responses harnessing the GALT. Here, we uncovered the epigenetic modifier MLL1 as a therapeutic target to promote IgA-producing PC. While our work used the model helminth *T. muris* to identify IgA regulation by MLL1, the need for promoting protective IgA through new-generation mucosal vaccines and treatments extends beyond helminth-specific vaccines (62), and is an area of intense current research. One benefit of histone modifiers as therapeutic targets is that their function can be inhibited by small molecule inhibitors (33, 63). Small molecule inhibitors of MLL1 are in development for treatment of MLL-rearranged acute lymphoblastic leukemias (64), thus in the future these inhibitors could be tested to examine whether they are able to promote IgA responses. It is important to note that the delivery of a MLL1 inhibitor would need to be B cell-targeted, given the roles of MLL1 in intestinal stem cells (28) and Th2 cells (27).

During an immune response, B cell differentiation is orchestrated by both extrinsic and intrinsic factors. The infecting pathogen or immunizing agent induces the activation of immune cells that will secrete cytokines and other factors into the microenvironment, creating a milieu that will influence the fate of B cells and type of antibody produced (52). These signals can then induce the function of transcription factors and epigenetic regulators within the nucleus, to promote or repress gene expression programs and ultimately, regulate B cell differentiation and function (65, 66). While some key inducers of IgA isotype switching have been discovered, such as RUNX3 and p300, TGFβ and retinoic acid (67–69), we reveal here that inhibiting MLL1 can significantly increase production of IgA^+^ PC, both *in vitro* and *in vivo*, and promotes the localization of CCR9^+^IgA^+^ cells in the colon lamina propria. There appeared to be a humoral response trade-off, however, as GC B cells were significantly reduced and showed diminished frequencies of IgG1^+^ cells. Nevertheless, identifying factors that can be targeted to promote IgA^+^ PC has the potential to be pursued in infections that rely on IgA for positive outcomes. Previous research has demonstrated the importance of IgA in protection against helminth and other gastrointestinal infections. For example, antigen-specific IgA was important for mounting protective responses against the model intestinal helminth parasite *Heligmosomoides polygyrus* by preventing worm development (70), and IgA was found to be critical for clearance and protection against rotavirus infection (71). Thus, identifying regulators of IgA in the GALT may lead to a better understanding of how to promote the production of protective IgA responses against different types of gastrointestinal infections.

These findings also point to non-overlapping roles of MLL1 and KMT2D (MLL2) in B cell biology, as *Kmt2d* deletion promotes GC B cell formation (31). Unique roles for these methyltransferases have also been noted in other systems, and is likely due to differing histone-modifying activity and gene targets (72–74). In contrast to KMT2D, we did not observe global H3K4me3 loss, but instead identified more discrete, site-specific modulation at approximately 300 gene loci. MLL1 has a wider array of histone-modifying activity and molecular partners, and thus future studies should delineate the exact histone modifications and target loci that are modified by MLL1 in GALT B cells.

In summary, our study has uncovered MLL1 as a regulator of GALT B cell responses. These results provide further understanding of the roles of histone modifiers in tailoring humoral immune responses. More broadly, improving our understanding on how to promote effective IgA-mediated responses in GALT will offer new avenues for oral and mucosal vaccine design.

## Acknowledgements

We thank Yu-Anne Yap, Moshe Olshansky, and members of the Good-Jacobson lab for technical assistance; Patricia Ernst for *Mll1* floxed mice, and staff of the Monash University FlowCore, Monash Animal Research Platform, Monash Histology Platform, and the Monash Genomics and Bioinformatics Platform.

## Author contributions

KLG-J conceived the study; KLG-J and YZ designed research; YZ, CC, AK, JP, AZ and RF performed research; KLG-J, CC, AK, DC and YZ analyzed data; CZ, DL and JRG provided intellectual input; CZ, LX, DLU, SM and JRG provided reagents and technical expertise; and KLG-J, YZ and CC wrote the manuscript.

## Abbreviations

CUT&Tag: Cleavage Under Targets and Tagmentation
GALT: Gut-associated lymphoid tissues
GC: germinal center
mLN: mesenteric lymph nodes
MLL1: mixed-lineage leukemia 1
PC: plasma cells

## References

1. Jourdan, P. M., P. H. L. Lamberton, A. Fenwick, and D. G. Addiss. 2018. Soil-transmitted helminth infections. Lancet 391: 252–265.

2. Zaini, A., K. L. Good-Jacobson, and C. Zaph. 2021. Context-dependent roles of B cells during intestinal helminth infection. PLoS Negl Trop Dis 15: e0009340.

3. WHO. Accessed 2 March 2020. Soil-transmitted helminth infections. *[online]* https://www.who.int/news-room/fact-sheets/detail/soil-transmitted-helminth-infections.

4. Crompton, D. W., and M. C. Nesheim. 2002. Nutritional impact of intestinal helminthiasis during the human life cycle. Annu Rev Nutr 22: 35–59.

5. Chelkeba, L., Z. Mekonnen, Y. Alemu, and D. Emana. 2020. Epidemiology of intestinal parasitic infections in preschool and school-aged Ethiopian children: a systematic review and meta-analysis. BMC Public Health 20: 117.

6. Pullan, R. L., J. L. Smith, R. Jasrasaria, and S. J. Brooker. 2014. Global numbers of infection and disease burden of soil transmitted helminth infections in 2010. Parasit Vectors 7: 37.

7. Dunn, J. C., H. C. Turner, A. Tun, and R. M. Anderson. 2016. Epidemiological surveys of, and research on, soil-transmitted helminths in Southeast Asia: a systematic review. Parasit Vectors 9: 31.

8. Zawawi, A., and K. J. Else. 2020. Soil-Transmitted Helminth Vaccines: Are We Getting Closer? Frontiers in immunology 11.

9. James, C. E., A. L. Hudson, and M. W. Davey. 2009. Drug resistance mechanisms in helminths: is it survival of the fittest? Trends Parasitol 25: 328–335.

10. Dunn, J. C., A. A. Bettis, N. Y. Wyine, A. M. M. Lwin, A. Tun, N. S. Maung, and R. M. Anderson. 2019. Soil-transmitted helminth reinfection four and six months after mass drug administration: results from the delta region of Myanmar. PLoS Negl Trop Dis 13: e0006591.

11. Dixon, H., C. E. Johnston, and K. J. Else. 2008. Antigen selection for future anti-Trichuris vaccines: a comparison of cytokine and antibody responses to larval and adult antigen in a primary infection. Parasite Immunol 30: 454–461.

12. Diemert, D. J., M. E. Bottazzi, J. Plieskatt, P. J. Hotez, and J. M. Bethony. 2018. Lessons along the Critical Path: Developing Vaccines against Human Helminths. Trends Parasitol 34: 747–758.

13. Grencis, R. K. 1993. Cytokine-mediated regulation of intestinal helminth infections: the Trichuris muris model. Ann Trop Med Parasitol 87: 643–647.

14. Bemark, M., M. J. Pitcher, C. Dionisi, and J. Spencer. 2024. Gut-associated lymphoid tissue: a microbiota-driven hub of B cell immunity. Trends Immunol 45: 211–223.

15. Piovesan, D., J. Tempany, A. Di Pietro, I. Baas, C. Yiannis, K. O’Donnell, Y. Chen, V. Peperzak, G. T. Belz, C. R. Mackay, G. K. Smyth, J. R. Groom, D. M. Tarlinton, and K. L. Good-Jacobson. 2017. c-Myb Regulates the T-Bet-Dependent Differentiation Program in B Cells to Coordinate Antibody Responses. Cell Rep 19: 461–470.

16. Spencer, J., and L. M. Sollid. 2016. The human intestinal B-cell response. Mucosal Immunology 9: 1113–1124.

17. Conrey, P. E., L. Denu, K. C. O’Boyle, I. Rozich, J. Green, J. Maslanka, J. B. Lubin, T. Duranova, B. L. Haltzman, L. Gianchetti, D. A. Oldridge, N. De Luna, L. A. Vella, D. Allman, J. M. Spergel, C. Tanes, K. Bittinger, S. E. Henrickson, and M. A. Silverman. 2023. IgA deficiency destabilizes homeostasis toward intestinal microbes and increases systemic immune dysregulation. Sci Immunol 8: eade2335.

18. Pabst, O. 2012. New concepts in the generation and functions of IgA. Nat Rev Immunol 12: 821–832.

19. Ramos, A. C. S., L. M. Oliveira, Y. Santos, M. C. S. Dantas, C. I. B. Walker, A. M. C. Faria, L. L. Bueno, S. S. Dolabella, and R. T. Fujiwara. 2022. The role of IgA in gastrointestinal helminthiasis: A systematic review. Immunol Lett 249: 12–22.

20. Fransen, F., E. Zagato, E. Mazzini, B. Fosso, C. Manzari, S. El Aidy, A. Chiavelli, A. M. D’Erchia, M. K. Sethi, O. Pabst, M. Marzano, S. Moretti, L. Romani, G. Penna, G. Pesole, and M. Rescigno. 2015. BALB/c and C57BL/6 Mice Differ in Polyreactive IgA Abundance, which Impacts the Generation of Antigen-Specific IgA and Microbiota Diversity. Immunity 43: 527–540.

21. Koyama, K., H. Tamauchi, M. Tomita, T. Kitajima, and Y. Ito. 1999. B-cell activation in the mesenteric lymph nodes of resistant BALB/c mice infected with the murine nematode parasite Trichuris muris. Parasitol Res 85: 194–199.

22. Clerc, M., G. Devevey, A. Fenton, and A. B. Pedersen. 2018. Antibodies and coinfection drive variation in nematode burdens in wild mice. Int J Parasitol 48: 785–792.

23. McGuire, C., W. C. Chan, and D. Wakelin. 2002. Nasal immunization with homogenate and peptide antigens induces protective immunity against Trichinella spiralis. Infect Immun 70: 7149–7152.

24. Yang, Y., Z. Zhang, J. Yang, X. Chen, S. Cui, and X. Zhu. 2010. Oral vaccination with Ts87 DNA vaccine delivered by attenuated Salmonella typhimurium elicits a protective immune response against Trichinella spiralis larval challenge. Vaccine 28: 2735–2742.

25. Roach, T. I., K. J. Else, D. Wakelin, D. J. McLaren, and R. K. Grencis. 1991. Trichuris muris: antigen recognition and transfer of immunity in mice by IgA monoclonal antibodies. Parasite Immunol 13: 1–12.

26. Gan, T., B. E. Li, B. P. Mishra, K. L. Jones, and P. Ernst. 2018. MLL1 Promotes IL-7 Responsiveness and Survival during B Cell Differentiation. J Immunol 200: 1682–1691.

27. Yamashita, M., K. Hirahara, R. Shinnakasu, H. Hosokawa, S. Norikane, M. Y. Kimura, A. Hasegawa, and T. Nakayama. 2006. Crucial Role of MLL for the Maintenance of Memory T Helper Type 2 Cell Responses. Immunity 24: 611–622.

28. Goveas, N., C. Waskow, K. Arndt, J. Heuberger, Q. Zhang, D. Alexopoulou, A. Dahl, W. Birchmeier, K. Anastassiadis, A. F. Stewart, and A. Kranz. 2021. MLL1 is required for maintenance of intestinal stem cells. PLoS Genet 17: e1009250.

29. Caganova, M., C. Carrisi, G. Varano, F. Mainoldi, F. Zanardi, P. L. Germain, L. George, F. Alberghini, L. Ferrarini, A. K. Talukder, M. Ponzoni, G. Testa, T. Nojima, C. Doglioni, D. Kitamura, K. M. Toellner, I. H. Su, and S. Casola. 2013. Germinal center dysregulation by histone methyltransferase EZH2 promotes lymphomagenesis. J Clin Invest 123: 5009–5022.

30. Béguelin, W., R. Popovic, M. Teater, Y. Jiang, Karen L. Bunting, M. Rosen, H. Shen, Shao N. Yang, L. Wang, T. Ezponda, E. Martinez-Garcia, H. Zhang, Y. Zheng, Sharad K. Verma, Michael T. McCabe, Heidi M. Ott, Glenn S. Van Aller, Ryan G. Kruger, Y. Liu, Charles F. McHugh, David W. Scott, Young R. Chung, N. Kelleher, R. Shaknovich, Caretha L. Creasy, Randy D. Gascoyne, K.-K. Wong, L. Cerchietti, Ross L. Levine, O. Abdel-Wahab, Jonathan D. Licht, O. Elemento, and Ari M. Melnick. 2013. EZH2 Is Required for Germinal Center Formation and Somatic EZH2 Mutations Promote Lymphoid Transformation. Cancer cell 23: 677–692.

31. Zhang, J., D. Dominguez-Sola, S. Hussein, J. E. Lee, A. B. Holmes, M. Bansal, S. Vlasevska, T. Mo, H. Tang, K. Basso, K. Ge, R. Dalla-Favera, and L. Pasqualucci. 2015. Disruption of KMT2D perturbs germinal center B cell development and promotes lymphomagenesis. Nat Med 21: 1190–1198.

32. Hatzi, K., H. Geng, A. S. Doane, C. Meydan, R. LaRiviere, M. Cardenas, C. Duy, H. Shen, M. N. C. Vidal, T. Baslan, H. P. Mohammad, R. G. Kruger, R. Shaknovich, A. M. Haberman, G. Inghirami, S. W. Lowe, and A. M. Melnick. 2019. Histone demethylase LSD1 is required for germinal center formation and BCL6-driven lymphomagenesis. Nat Immunol 20: 86–96.

33. Di Pietro, A., J. Polmear, L. Cooper, T. Damelang, T. Hussain, L. Hailes, K. O’Donnell, V. Udupa, T. Mi, S. Preston, A. Shtewe, U. Hershberg, S. J. Turner, N. L. La Gruta, A. W. Chung, D. M. Tarlinton, C. D. Scharer, and K. L. Good-Jacobson. 2022. Targeting BMI-1 in B cells restores effective humoral immune responses and controls chronic viral infection. Nat Immunol 23: 86–98.

34. Good-Jacobson, K. L., Y. Chen, A. K. Voss, G. K. Smyth, T. Thomas, and D. Tarlinton. 2014. Regulation of germinal center responses and B-cell memory by the chromatin modifier MOZ. Proc Natl Acad Sci U S A 111: 9585–9590.

35. Kealy, L., A. Di Pietro, L. Hailes, S. Scheer, L. Dalit, J. R. Groom, C. Zaph, and K. L. Good-Jacobson. 2020. The Histone Methyltransferase DOT1L Is Essential for Humoral Immune Responses. Cell Rep 33: 108504.

36. Nakata, Y., A. C. Brignier, S. Jin, Y. Shen, S. I. Rudnick, M. Sugita, and A. M. Gewirtz. 2010. c-Myb, Menin, GATA-3, and MLL form a dynamic transcription complex that plays a pivotal role in human T helper type 2 cell development. Blood 116: 1280–1290.

37. Kwon, K., C. Hutter, Q. Sun, I. Bilic, C. Cobaleda, S. Malin, and M. Busslinger. 2008. Instructive role of the transcription factor E2A in early B lymphopoiesis and germinal center B cell development. Immunity 28: 751–762.

38. Jude, C. D., L. Climer, D. Xu, E. Artinger, J. K. Fisher, and P. Ernst. 2007. Unique and independent roles for MLL in adult hematopoietic stem cells and progenitors. Cell Stem Cell 1: 324–337.

39. Antignano, F., S. C. Mullaly, K. Burrows, and C. Zaph. 2011. Trichuris muris infection: a model of type 2 immunity and inflammation in the gut. J Vis Exp.

40. Livak, K. J., and T. D. Schmittgen. 2001. Analysis of relative gene expression data using real-time quantitative PCR and the 2(-Delta Delta C(T)) Method. Methods 25: 402–408.

41. Watt, K. A., D. H. Nussey, R. Maclellan, J. G. Pilkington, and T. N. McNeilly. 2016. Fecal antibody levels as a noninvasive method for measuring immunity to gastrointestinal nematodes in ecological studies. Ecol Evol 6: 56–67.

42. Smeekens, J. M., B. T. Johnson-Weaver, A. L. Hinton, M. A. Azcarate-Peril, T. P. Moran, R. M. Immormino, J. R. Kesselring, E. C. Steinbach, K. A. Orgel, H. F. Staats, A. W. Burks, P. J. Mucha, M. T. Ferris, and M. D. Kulis. 2020. Fecal IgA, Antigen Absorption, and Gut Microbiome Composition Are Associated With Food Antigen Sensitization in Genetically Susceptible Mice. Front Immunol 11: 599637.

43. Kaya-Okur, H. S., S. J. Wu, C. A. Codomo, E. S. Pledger, T. D. Bryson, J. G. Henikoff, K. Ahmad, and S. Henikoff. 2019. CUT&Tag for efficient epigenomic profiling of small samples and single cells. Nat Commun 10: 1930.

44. Cheshire, C., charlotte-west, T. Ronkko, n.-c. bot, H. Patel, tamara-hodgetts, D. Ladd, A. Thiery, C. Fields, J. Deu-Pons, P. Ewels, S. Moller, and K. Menden. 2024. nf-core/cutandrun: nf-core/cutandrun v3.2.2 Iridium Ibis (3.2.2). Zenodo 10.5281/zenodo.10606804.

45. Zhang, Y., T. Liu, C. A. Meyer, J. Eeckhoute, D. S. Johnson, B. E. Bernstein, C. Nusbaum, R. M. Myers, M. Brown, W. Li, and X. S. Liu. 2008. Model-based analysis of ChIP-Seq (MACS). Genome Biol 9: R137.

46. Heinz, S., C. Benner, N. Spann, E. Bertolino, Y. C. Lin, P. Laslo, J. X. Cheng, C. Murre, H. Singh, and C. K. Glass. 2010. Simple combinations of lineage-determining transcription factors prime cis-regulatory elements required for macrophage and B cell identities. Mol Cell 38: 576–589.

47. Liao, Y., G. K. Smyth, and W. Shi. 2014. featureCounts: an efficient general purpose program for assigning sequence reads to genomic features. Bioinformatics 30: 923–930.

48. Liao, Y., G. K. Smyth, and W. Shi. 2019. The R package Rsubread is easier, faster, cheaper and better for alignment and quantification of RNA sequencing reads. Nucleic Acids Res 47: e47.

49. Law, C. W., Y. Chen, W. Shi, and G. K. Smyth. 2014. voom: Precision weights unlock linear model analysis tools for RNA-seq read counts. Genome Biol 15: R29.

50. Ritchie, M. E., B. Phipson, D. Wu, Y. Hu, C. W. Law, W. Shi, and G. K. Smyth. 2015. limma powers differential expression analyses for RNA-sequencing and microarray studies. Nucleic Acids Res 43: e47.

51. Zaini, A., L. Dalit, A. A. Sheikh, Y. Zhang, D. Thiele, J. Runting, G. Rodrigues, J. Ng, M. Bramhall, S. Scheer, L. Hailes, J. R. Groom, K. L. *Good-Jacobson, C. *Zaph, and C.-s. authors. 2023. Heterogeneous Tfh cell populations that develop during enteric helminth infection predict the quality of type 2 protective response. Mucosal Immunol 16: 642–657.

52. Good-Jacobson, K. L. 2018. Strength in diversity: Phenotypic, functional, and molecular heterogeneity within the memory B cell repertoire. Immunol Rev 284: 67–78.

53. Montaño, K. J., C. Cuéllar, and J. Sotillo. 2021. Rodent Models for the Study of Soil-Transmitted Helminths: A Proteomics Approach. Front Cell Infect Microbiol 11: 639573.

54. Blackwell, N. M., and K. J. Else. 2001. B cells and antibodies are required for resistance to the parasitic gastrointestinal nematode Trichuris muris. Infect Immun 69: 3860–3868.

55. Oeser, K., C. Schwartz, and D. Voehringer. 2015. Conditional IL-4/IL-13-deficient mice reveal a critical role of innate immune cells for protective immunity against gastrointestinal helminths. Mucosal Immunol 8: 672–682.

56. Turner, J. D., H. Faulkner, J. Kamgno, F. Cormont, J. Van Snick, K. J. Else, R. K. Grencis, J. M. Behnke, M. Boussinesq, and J. E. Bradley. 2003. Th2 cytokines are associated with reduced worm burdens in a human intestinal helminth infection. J Infect Dis 188: 1768–1775.

57. Reboldi, A., and J. G. Cyster. 2016. Peyer’s patches: organizing B-cell responses at the intestinal frontier. Immunol Rev 271: 230–245.

58. Pabst, O., L. Ohl, M. Wendland, M. A. Wurbel, E. Kremmer, B. Malissen, and R. Forster. 2004. Chemokine receptor CCR9 contributes to the localization of plasma cells to the small intestine. J Exp Med 199: 411–416.

59. Hayes, K. S., and R. K. Grencis. 2021. Trichuris muris and comorbidities - within a mouse model context. Parasitology 148: 1–9.

60. Kamada, N., K. Sakamoto, S. U. Seo, M. Y. Zeng, Y. G. Kim, M. Cascalho, B. A. Vallance, J. L. Puente, and G. Nunez. 2015. Humoral Immunity in the Gut Selectively Targets Phenotypically Virulent Attaching-and-Effacing Bacteria for Intraluminal Elimination. Cell Host Microbe 17: 617–627.

61. Dou, Y., T. A. Milne, A. J. Ruthenburg, S. Lee, J. W. Lee, G. L. Verdine, C. D. Allis, and R. G. Roeder. 2006. Regulation of MLL1 H3K4 methyltransferase activity by its core components. Nat Struct Mol Biol 13: 713–719.

62. Oh, J. E., E. Song, M. Moriyama, P. Wong, S. Zhang, R. Jiang, S. Strohmeier, S. H. Kleinstein, F. Krammer, and A. Iwasaki. 2021. Intranasal priming induces local lung-resident B cell populations that secrete protective mucosal antiviral IgA. Sci Immunol 6: eabj5129.

63. Polmear, J., L. Hailes, M. Olshansky, M. Rischmueller, E. L’Estrange-Stranieri, A. L. Fletcher, M. L. Hibbs, V. L. Bryant, and K. L. Good-Jacobson. 2023. Targeting BMI-1 to deplete antibody-secreting cells in autoimmunity. Clin Transl Immunology 12: e1470.

64. Chern, T. R., L. Liu, E. Petrunak, J. A. Stuckey, M. Wang, D. Bernard, H. Zhou, S. Lee, Y. Dou, and S. Wang. 2020. Discovery of Potent Small-Molecule Inhibitors of MLL Methyltransferase. ACS Med Chem Lett 11: 1348–1352.

65. Zhang, Y., and K. L. Good-Jacobson. 2019. Epigenetic regulation of B cell fate and function during an immune response. Immunol Rev 288: 75–84.

66. Cooper, L., H. Xu, J. Polmear, L. Kealy, C. Szeto, E. S. Pang, M. Gupta, A. Kirn, J. J. Taylor, K. J. L. Jackson, B. J. Broomfield, A. Nguyen, C. G. da Graça, N. L. La Gruta, D. T. Utzschneider, J. R. Groom, L. Martelotto, I. A. Parish, M. O’Keeffe, C. D. Scharer, S. Gras, and K. L. Good-Jacobson. 2024. Type I interferons induce an epigenetically distinct memory B cell subset in chronic viral infection. Immunity 57: 1037–1055.

67. Park, S. R., E. K. Lee, B. C. Kim, and P. H. Kim. 2003. p300 cooperates with Smad3/4 and Runx3 in TGFbeta1-induced IgA isotype expression. Eur J Immunol 33: 3386–3392.

68. Cerutti, A. 2008. The regulation of IgA class switching. Nat Rev Immunol 8: 421–434.

69. Reboldi, A., T. I. Arnon, L. B. Rodda, A. Atakilit, D. Sheppard, and J. G. Cyster. 2016. IgA production requires B cell interaction with subepithelial dendritic cells in Peyer’s patches. Science 352: aaf4822.

70. McCoy, K. D., M. Stoel, R. Stettler, P. Merky, K. Fink, B. M. Senn, C. Schaer, J. Massacand, B. Odermatt, H. C. Oettgen, R. M. Zinkernagel, N. A. Bos, H. Hengartner, A. J. Macpherson, and N. L. Harris. 2008. Polyclonal and specific antibodies mediate protective immunity against enteric helminth infection. Cell Host Microbe 4: 362–373.

71. Blutt, S. E., A. D. Miller, S. L. Salmon, D. W. Metzger, and M. E. Conner. 2012. IgA is important for clearance and critical for protection from rotavirus infection. Mucosal Immunol 5: 712–719.

72. Wang, P., C. Lin, E. R. Smith, H. Guo, B. W. Sanderson, M. Wu, M. Gogol, T. Alexander, C. Seidel, L. M. Wiedemann, K. Ge, R. Krumlauf, and A. Shilatifard. 2009. Global analysis of H3K4 methylation defines MLL family member targets and points to a role for MLL1-mediated H3K4 methylation in the regulation of transcriptional initiation by RNA polymerase II. Mol Cell Biol 29: 6074–6085.

73. Mishra, B. P., K. M. Zaffuto, E. L. Artinger, T. Org, H. K. Mikkola, C. Cheng, M. Djabali, and P. Ernst. 2014. The histone methyltransferase activity of MLL1 is dispensable for hematopoiesis and leukemogenesis. Cell Rep 7: 1239–1247.

74. Huang, X., J. Yan, M. Zhang, Y. Wang, Y. Chen, X. Fu, R. Wei, X. L. Zheng, Z. Liu, X. Zhang, H. Yang, B. Hao, Y. Y. Shen, Y. Su, X. Cong, M. Huang, M. Tan, J. Ding, and M. Geng. 2018. Targeting Epigenetic Crosstalk as a Therapeutic Strategy for EZH2-Aberrant Solid Tumors. Cell 175: 186–199 e119.

